# Ketamine Effects on Energy Metabolism, Functional Connectivity and Working Memory in Healthy Humans

**DOI:** 10.1101/2023.02.21.529425

**Authors:** Naomi R. Driesen, Peter Herman, Margaret A. Rowland, Garth Thompson, Maolin Qiu, George He, Sarah Fineberg, Daniel S. Barron, Lars Helgeson, Cheryl Lacadie, Robert Chow, Ralitza Gueorguieva, Teo-Carlo Straun, John H. Krystal, Fahmeed Hyder

**Author notes:** Drs. Krystal and Hyder provided complementary, equivalent contributions to this paper.

## Abstract

Working memory (WM) is a crucial resource for temporary memory storage and the guiding of ongoing behavior. N-methyl-D-aspartate glutamate receptors (NMDARs) are thought to support the neural underpinnings of WM. Ketamine is an NMDAR antagonist that has cognitive and behavioral effects at subanesthetic doses. To shed light on subanesthetic ketamine effects on brain function, we employed a multimodal imaging design, combining gas-free calibrated functional magnetic resonance imaging (fMRI) measurement of oxidative metabolism (CMRO_2_), resting-state cortical functional connectivity assessed with fMRI, and WM-related fMRI. Healthy subjects participated in two scan sessions in a randomized, double-blind, placebo-controlled design. Ketamine increased CMRO_2_ and cerebral blood flow (CBF) in prefrontal cortex (PFC) and other cortical regions. However, resting-state cortical functional connectivity was not affected. Ketamine did not alter CBF-CMRO_2_ coupling brain-wide. Higher levels of basal CMRO_2_ were associated with lower task-related PFC activation and WM accuracy impairment under both saline and ketamine conditions. These observations suggest that CMRO_2_ and resting-state functional connectivity index distinct dimensions of neural activity. Ketamine’s impairment of WM-related neural activity and performance appears to be related to its ability to produce cortical metabolic activation. This work illustrates the utility of direct measurement of CMRO_2_ via calibrated fMRI in studies of drugs that potentially affect neurovascular and neurometabolic coupling.

## Introduction

Preclinical research, computational modeling, and human neuroimaging studies highlight a critical role for N-methyl-D-aspartate glutamate receptors (NMDARs) in working memory (WM). WM is a key cognitive resource that provides temporary memory storage and guidance of ongoing behavior. The neural activity underlying WM depends on the recurrent excitation of pyramidal cells in layer III of lateral prefrontal cortex (PFC) (1–6). Computational models suggest that NMDARs are vital for this recurrent excitation in WM (7–10). This hypothesis has been corroborated *in vivo* in the non-human primate (11). In humans, the NMDAR antagonist ketamine reduces task-related activation in the functional magnet resonance imaging (fMRI) blood-oxygenation level dependent (BOLD) signal during WM (12–14), and impairs WM-related performance (7, 12–15), albeit with considerable inter-subject variability in these findings (e.g., 13).

NMDAR antagonists reduce the incremental WM-related cortical activation in the context of increased resting (baseline) cortical activity. In animals, administration of NMDAR antagonists reduce GABA neuronal function (16, 17) and reductions in inhibition contribute to increased glutamate release (18), thereby raising the rate of resting cortical activity (19). In non-human primate PFC, while WM “delay” pyramidal neurons in layer III are inhibited by local ketamine administration, the pyramidal neurons in layer V that reflect WM response are activated by systemic ketamine (11). In humans, intravenous administration of ketamine increases resting brain activity as measured by positron emission tomography (PET) (20–23), resting fMRI functional connectivity (24), an MRS-based measure of glutamate release (25), and a PET indirect estimate of glutamate release (26–28). Thus, while local administration of NMDAR antagonists reduces synaptic activation, systemic administration disinhibits the cortex. Cortical disinhibition by ketamine could directly impair WM by reducing cortical tuning (7, 29, 30). It is also possible that cortical activation recruits interneurons to produce inhibition of WM-related function, i.e., the “downside” of the “inverted-U curve” describing the relationship between cortical activation and WM function (31, 32).

The current study evaluated the relationship between ketamine effects on baseline and WM-related cortical activity using a calibrated fMRI (CALIB) approach (33–35). Traditional fMRI is poorly suited to testing this hypothesis since ketamine, like many drugs, may disturb neurovascular and neurometabolic coupling (36). BOLD-fMRI signals reflect the confluence of changes in cerebral blood flow (CBF), cerebral blood volume (CBV), and cerebral metabolism (CMRO_2_) (37). Through a biophysical model based on decades of animal and now human experimentation (35, 38–41), CALIB helps assess changes in CMRO_2_ disassociated from those in CBF or CBV and the BOLD signal, thus facilitating interpretation of neural activity changes between states (31, 32).

Human studies must measure the dynamic range of the BOLD signal in a given voxel (M) for CALIB. Typically, M is determined by varying CO_2_ (and/or O_2_) levels via gas inhalation (38, 42–44). M-values obtained by these studies show variations across laboratories, brain regions, and magnetic field strengths (39). Furthermore, the experimental data obtained during gas exposures represent a very small pseudo-linear portion of a highly non-linear (i.e., asymptotic) relationship between very large CBF changes in relation to much smaller BOLD signal changes. During gas exposure, the measured BOLD signal increases are 2-3% and CBF increases are 50-100% (33, 45), whereas the exponential fitting that defines the M-value from the asymptotic limit is reached when CBF increases by several times higher (34, 46). Moreover, it is difficult to reach large CBF changes in humans without inducing adverse cognitive effects and changes in neural or metabolic activities (47–51).

This is one of the first human studies to employ a novel, gas-free approach to M assessment. This non-gaseous CALIB integrates the BOLD-sensitive transverse relaxation rate information from spin-echo (R_2_) and gradient-echo (R_2_*) BOLD imaging (i.e., R_2_’= R_2_*-R_2_) as well as CBF measured by arterial spin labeling (ASL) to achieve absolute quantification of CMRO_2_ (35). Specifically, measurements of CBF and R_2_’ are combined in a biophysical mathematical model to assess CMRO_2_. Note that the product of R_2_’ and echo time (TE) is the same M as theoretically derived from gas exposure CALIB experiments (37, 38). CMRO_2_ measures derived by this method are nearly identical to direct measurements with ^13^C MRS (35) and these metabolic measures are comparable to electrical recordings of neuronal firing (35, 52).

In this study, we leveraged CALIB to assess the impact of ketamine on resting- and WM-related cortical activity, enabling the exploration of the relationship between cortical disinhibition and WM-related neural activity and performance. To accomplish this, we employed a multimodal imaging design, combining task-related fMRI activation in PFC with behavioral performance, resting-state fMRI functional connectivity, and non-gaseous CALIB to estimate CMRO_2_ and CBF. We tested healthy subjects in two different scan sessions in a randomized, double-blind, placebo-controlled design. All behavioral and fMRI metrics were measured during saline and during ketamine. On each test day, we assessed, using interleaved scans, CBF/CMRO_2_ and functional connectivity during rest as well as task-related brain activation during a spatial WM task. We previously showed that ketamine reduces task-related activation using the WM task employed in this study (12–14). We hypothesized that ketamine would increase CMRO_2_ in the PFC, and that these increases would be associated with decreases in WM accuracy and task-related activation. We also expected that ketamine-associated increases in CMRO_2_ and functional connectivity would parallel each other, as we have previously shown that ketamine increases functional connectivity as quantified with Global Brain Connectivity (GBC). We found that ketamine increased prefrontal CMRO_2_ and CBF and that CBF and CMRO_2_ couplings during saline and during ketamine were very tight. The differential impact of ketamine (i.e., ketamine – saline) on resting cortical activity as measured by CMRO_2_ or CBF did not predict the impairment of WM-related cortical activation or task performance. Rather, the CMRO_2_ or CBF values obtained during ketamine or saline (single state measurement) predicted both task-related cortical activation and WM accuracy. The current findings suggest that the multimodal design incorporating behavior with various fMRI methods is needed for non-invasive assessment of the neural effects of drugs that impact neurovascular and neurometabolic coupling.

## Materials and Methods

### Participants

Twenty-three healthy volunteers participated in this randomized, counterbalanced, double-blinded study. Every participant had two fMRI scanning sessions which lasted two hours and occurred at least two weeks apart. During each session, the volunteers were infused either with saline or a subanesthetic dose of ketamine. Prior to the study onset, participants received written explanations of the study and potential side effects. All participants gave their written consent to participate in the study approved by the Yale University Human Investigation Committee. The experiments were performed at the Yale Magnetic Resonance Research Center on a 3T Siemens Prisma scanner using 64-channel head/neck coil.

On the morning of the scan, participants were assessed with the Positive and Negative Syndrome Scale (PANSS; 53) and the Clinician-Administered Dissociative States Scale (CADSS; 54). They also received training on the spatial WM task (55). Two intravenous catheters were placed: one for infusing saline or ketamine, and another for drawing blood samples. Each subject’s pulse, respiration, oxygen saturation, and heart rate were monitored and recorded throughout the scanning session using BioPac model MP150 data acquisition system (BioPac Systems, Inc., Goleta, CA, USA). For safety purposes, vitals were also monitored using the Invivo Millenia 3155MVS system (Invivo Research, Inc., Orlando, FL, USA). Ketamine was infused in the MRI scanner starting with a bolus of 0.23 mg/kg/minute ketamine which was subsequently reduced to 0.58 mg/kg/hour ketamine for 30 minutes. After 30 minutes, the infusion was reduced further to 0.29 mg/kg/hour to yield steady plasma ketamine levels throughout the session. Saline infusion followed the same protocol. During the session, four blood samples were collected to measure plasma ketamine level (56). The PANSS and the CADSS were administered immediately after the scan to capture experiences during ketamine administration.

### Spatial Working Memory Task

During specified runs, subjects performed the spatial WM task. Stimuli were projected onto a screen that subjects viewed via a mirror. For the hard (4-item) task, stimuli were four red solid circles presented sequentially in locations ranging from ±3.6° to ±11.5° from the center. Subjects fixated on a dot at the center of the visual field for 3,250 ms. Each solid red circle was presented for 1 s with a 250 ms inter-stimulus interval. After a 13 s retention interval (RI), a ring, serving as a probe, appeared on the screen for 1 s. Subjects pressed a button to indicate whether the probe location matched one of the previously presented circles. After subjects responded to the probe, the fixation dot changed to a cross, which became a dot at the start of the next trial. The inter-trial interval (ITI) was 14 s. The easy (2-item) task was the same as the 4-item except only two circles were presented. An additional 2,500 ms was added to the ITI to make the 4-item and 2-item trials the same length. The control (no WM) task closely resembled the hard task except that the color of the circles was grey. Participants were instructed to simply attend to the stimuli without attempting to remember their locations. They received all types of trials in a fixed, pseudo-random order.

### MRI data acquisitions

During the two-hour session, we collected multimodal images for calculating CMRO_2_ and for assessing resting functional connectivity and task-related activation to the spatial WM tasks. Scan parameters and order are detailed in Table 1. First, we acquired T1-weighted MPRAGE structural images in a sagittal orientation and T1 FLASH images. The T1 FLASH images were co-planar to the images used in the calibrated fMRI. For the calibrated fMRI, we collected T2 and T2* maps with different echo times. After this, we acquired three pulsed arterial spin labeling (PASL) echo planar imaging (EPI) scans. Before and after the PASL scans, we acquired proton density (PD) scans with similar parameters but with disabled arterial labeling. In alternation with the PASL scans, we acquired six spatial WM EPI scans. For registration purposes, we collected T1 FLASH images co-planar with the WM fMRI scans at the middle and end of the scanning session. We also acquired resting-state multiband fMRI single-shot gradient-echo EPI scans in two successive runs. Additionally, we collected T1 FLASH images co-planar with the resting-state scans.

**Table 1.** Key Scan Parameters. Note – Co-planars are structural scans completed for the purpose of coregistration, TA = time of acquisition, TE = time to echo.

| Scan Type | TA | TE | TR | Flip Angle (Degrees) | Voxel Size (mm) | Slices | Matrix |
| --- | --- | --- | --- | --- | --- | --- | --- |
| 3 plane localizer | 0:10 | 5 | 14 | 40 | 1.1 x 1.1 x 5.0 | 5 | 280 x 280 |
| Three Dimensional MPRAGE (High Quality Structural Scan) | 5:12 | 2.98 | 2300 | 9 | 1.0 x 1.0 x 1.0 | 176 | 320 x 320 |
| Axial Oblique T1 flash (co-planar for ASL) | 2:35 | ----- | 300 | 60 | 1.0 x 1.0 x 5.0 | 20 | 256 x 256 |
| Proton Density | 1:40 | 26 | 10000 | 90 | 4.0 x 4.0 x 5.0 | 20 | 256 x 256 |
| T2* Mapping, gradient echo | 0:20 | 23, 30, 50, 80, 120, 170, 250 | 20000 | 90 | 4.0 x 4.0 x 5.0 | 20 | 256 x 256 |
| T2 Mapping, spin echo | 0:20 | 41, 70, 110, 160, 220, 300, 400 | 20000 | 90 | 4.0 x 4.0 x 5.0 | 20 | 256 x 256 |
| N-back (conventional BOLD) | 4:17 | 30 | 1500 | 80 | 4.0 x 4.0 x 4.0 | 27 | 256 x 256 |
| ASL | 5:09 | 26 | 3000 | 90 | 4.0 x 4.0 x 5.0 | 20 | 256 x 256 |
| N-back (conventional BOLD) | 4:17 | 30 | 1500 | 80 | 4.0 x 4.0 x 4.0 | 27 | 256 x 256 |
| ASL | 5:09 | 26 | 3000 | 90 | 4.0 x 4.0 x 5.0 | 20 | 256 x 256 |
| N-back (conventional BOLD) | 4:17 | 30 | 1500 | 80 | 4.0 x 4.0 x 4.0 | 27 | 256 x 256 |
| Multiband Rest | 4:15 | 30 | 1000 | 55 | 2.0 x 2.0 x 2.0 | 75 | 220 x 220 |
| N-back (conventional BOLD) | 4:17 | 30 | 1500 | 80 | 4.0 x 4.0 x 4.0 | 27 | 256 x 256 |
| ASL | 5:09 | 26 | 3000 | 90 | 4.0 x 4.0 x 5.0 | 20 | 256 x 256 |
| N-back (conventional BOLD) | 4:17 | 30 | 1500 | 80 | 4.0 x 4.0 x 4.0 | 27 | 256 x 256 |
| T1 Co-Planar for N-back | 2:35 | 2.43 | 300 | 60 | 1.0 x 1.0 x 4.0 | 27 | 256 x 256 |
| T1 Coplanar for Multiband Rest | 2:39 | 2.61 | 440 | 70 | 0.9 x 0.9 x 2.0 | 75 | 220 x 220 |
| Proton Density | 1:40 | 26 | 10000 | 90 | 4.0 x 4.0 x 5.0 | 20 | 256 x 256 |
| Multiband Rest | 4:15 | 30 | 1000 | 55 | 2.0 x 2.0 x 2.0 | 75 | 220 x 220 |
| N-back (conventional BOLD) | 4:17 | 30 | 1500 | 80 | 4.0 x 4.0 x 4.0 | 27 | 256 x 256 |
| Multiband Rest | 4:15 | 30 | 1000 | 55 | 2.0 x 2.0 x 2.0 | 75 | 220 x 220 |

### Gas-free calibrated fMRI

The BOLD signal is an indirect indicator of the neural activity. But BOLD signal is a composite signal which depends on hemodynamic (CBV and CBF) and metabolic changes (CMRO_2_) in the brain. While the hemodynamic signals change indirectly with the neural activity, the metabolic signal is a direct representation of neural activity since the major origin of brain energy is the oxidative phosphorylation (57). The goal of calibrated fMRI is to extract CMRO_2_ signal from BOLD signal, using CBV and CBF signal, but along with additional adjustments. Because CBF and CBV are dependent on each other, they have a power law relationship (58) (CBV = k · CBF^α^, where k is a constant), which is defined by the constant α, the Grubb-coefficient, which ranges from 0.1-0.3 in rat and human brains (42, 59). Therefore, CMRO_2_ change (ΔCMRO_2_) between two conditions (C1 and C2, e.g., for saline or ketamine, respectively) can be calculated from simultaneously measured changes in BOLD and CBF with the following equation:

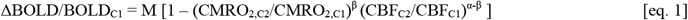

Parameters α and β are relatively stable and uniform throughout the brain and can be considered as region and condition independent (35, 38, 60). The constant β defines the power law dependence between the BOLD signal and deoxyhemoglobin concentration (38, 43). Parameter M is the maximum possible BOLD change without metabolic changes, which is frequently defined by CO_2_ inhalation (the gas-based BOLD calibration). However, M, an echo-time (TE) dependent variable, can be measured by defining the reversible BOLD relaxation component (R2’), which incorporates the resting CMRO_2_ and CBF conditions for a subject (38, 43). In eq. 1, M (the product of TE and R2’) is the unknown parameter which must be independently measured in each subject so the CMRO_2_ can be calculated. For a detailed explanation of the biophysical model, please see Supplemental Information (SI), Methods.

### Motion Correction

Motion correction was completed across all modalities via a common standard. CALIB and task/resting fMRI scans were motion corrected, and volumes with motion >2 mm in the z, y, or z direction or >3 degrees of pitch, yaw, or roll were identified. If out-of-range volumes were near the beginning or end of the scan, those volumes were eliminated. Otherwise, the entire scan was deleted.

### CBF and CMRO_2_ Image analysis

CMRO_2_ and CBF Images were computed via a pipeline using code from MATLAB, Analysis of Functional NeuroImages (AFNI), Bioimage Suite, and FMRIB Software Library (FSL). Images were converted to Neuroimaging Informatics Technology Initiative format (NIFTI), and brain extraction was performed on structural images using FSL (61). Scans were registered to the individual structural scan using the BioImage Suite software package (https://medicine.yale.edu/bioimaging/suite/). Using the Voxel-based Morphometry Toolbox (62), we created a grey matter (GM) mask for each subject at 70% probability threshold.

For the CBF measurement, we used PASL acquisitions to measure tissue perfusion (63). We calculated the absolute CBF images from the PASL and PD scans. First, we calculated the mean PD and the tagged/untagged time series were spatially filtered with a Gaussian filter (FWHM = [6 6 1] mm in the third direction). Untagged and tagged PASL images were separately realigned to the middle image of the series with AFNI (64, 65) and motion corrected. After motion correction, tagged and untagged images were time shifted using the sine interpolation method (66). The tagged and untagged volume series were co-registered to the mean PD image. Then, implementing QUantitative Imaging of Perfusion using a Single Subtraction method (QUIPSS-II; 67, 68), we calculated the absolute CBF in native space. CBF values from the three different runs during the fMRI session were averaged. To avoid artefacts, we removed voxels from further analysis of the CBF series whose values were below 5 or above 600 ml/min/100g. For the equations and parameters of QUIPSS-II, please see SI, Methods.

The T_2_* and T_2_ maps were calculated per voxel using single exponential model fitting to the multi-TE data in the native space. The T_2_ and T_2_* EPI scans were spatially realigned to the average PD image. Quantified T_2_ values only in the range from 5 ms to 150 ms and T_2_* values in the range from 5 ms to 100 ms were considered for further processing to avoid artifactual R_2_ and R_2_* values. R_2_ and R_2_* maps were calculated as 1/T_2_ and 1/T_2_*, respectively. The R_2_’ maps were the difference between R_2_* and R_2_ maps (SI, eq. 2.3) and thresholded to values above zero. Those voxels which were outside the CBF and R2s criteria were not used in the final analysis.

The R_2_’ and CBF maps were normalized to Montreal Neurological Institute (MNI) space and the CMRO_2_ changes were calculated (SI, eq. 5.2) with parameters α = 0.2 (42, 59), β = 1.5 (69, 70). To relate changes between conditions, we computed relative changes in CBF (rCBF = (CBFC1 - CBFC2) / CBFC2) and CMRO_2_ (rCMRO_2_ = (CMRO_2_,C1 – CMRO_2_,C2) / CMRO_2_,C2). Absolute CBF was calculated from PASL signal using QUIPSS-II in every condition. Absolute CMRO_2_ calculation, while it is possible by SI eq. 4.3, requires the estimation of the unmeasured A and k parameters. While some groups are claiming this approach for absolute CMRO_2_ by CALIB (71, 72), we prefer to use relative changes in CMRO_2_. However, based on preliminary data analysis, we could estimate A = 4 and k = 0.8, since these parameters move CMRO_2_ data values into the expected range as measured by PET data (57).

### Region of Interest Analysis

Whole-brain R_2_’, CBF, and CMRO_2_ maps were segmented into regions of interest (ROIs) after the application of individual GM masks (SI Figure S1). The GM masks were segmented from individual high resolution MPRAGE images using the Voxel-Based Morphology toolbox (http://dbm.neuro.uni-jena.de/vbm/). The selected ROIs were the standard Brodmann areas (BA) based on the Yale Brodmann ObjectMap (73). Additionally, we selected three regions for statistical comparisons: PFC, somatosensory and motor cortex (SSM), and visual cortex (VC). PFC was defined by combining BA 8, 9, 10, 11, 32, 44, 45, 46, and 47; SSM from BA 1, 4, 5, and 6; VC from BA 17, 18 and 19. Dorsolateral Prefrontal Cortex (DLPFC) was created from BA 9 and BA 46. Left and right regions were not separated. To minimize the influence of outliers and non-normal distributions, the median value for the intersection of BA regions and the individual GM matter mask was calculated and those values entered into statistical analysis.

### Resting State Functional Connectivity

The high resolution multiband BOLD data was used for analyzing functional connectivity via GBC. We computed the correlation between each voxel and all other voxels in the brain, returning an average correlation. Our primary analysis was with global signal. We also computed GBC without global signal. Detailed information regarding the GBC analyses is in SI, Methods. ROIs were computed in the same fashion as for the R2’, CBF, and CMRO_2_ analysis above.

### Task-Related Activation

Images were corrected to the middle image of the run for motion using MCFLIRT (61), corrected for slice-timing using Fourier-space time-series phase-shifting, smoothed with a Gaussian kernel of FWHM 5 mm and normalized with a single multiplicative factor. Temporal auto-correlation was estimated and corrected using FMRIB’s Improved Linear Model (FILM) (74). Images were high pass filtered (f > 0.01 Hz). Registration of all images was achieved by co-registering each individual’s co-planar image to their high-resolution structural scan. Then, the high-resolution structural scan was co-registered to the MNI standard brain supplied with FSL. To obtain nuisance regressors for cerebral spinal fluid (CSF) and white matter, images were segmented using the Voxel-Based Morphology toolbox (http://dbm.neuro.uni-jena.de/vbm/) into CSF, GM, and white matter. Average timecourses for CSF and white matter were then derived.

### Computation of Timecourses

First-level analyses were performed with motion parameters as well as CSF and white matter timecourses added to the regression. In addition, we added regressors for pulse and respiration. The residual image, i.e., the brain image with the influence of motion, CSF, white matter, and physiological variables removed, was entered into an ROI analysis. ROI analysis was completed using software tools from the Biomedical Informatics Research Network, a NIH/NHCRR consortium of university imaging centers (http://www.nitrc.org/projects/bxh_xcede_tools/). Scans with motion > 2 mm in the z, y, or z direction or > 3 degrees of pitch, yaw, or roll were eliminated unless the excess motion was confined to a few images that could simply be omitted by deleting these trials from the analysis. We used anatomical ROIs with a 10 mm radius previously developed for this task. The average timecourse for each ROI and trial type was then extracted from the pre-processed data for each scan. Each time point was expressed as a percent signal change from the pre-stimulus baseline, which consisted of three images collected at the beginning of the fixation period. At this stage of the analysis, information about percent signal change at each time point, i.e., for each repetition time (TR), ROI, and trial type, was maintained. Time points were averaged to early retention (8.75-17.75 s), later retention (18-24 s) and response (27-34.5 s) phases. To derive a variable reflecting task-related network activation, we combined the ROIs for inferior frontal gyrus (BA 45), anterior corpus callosum (BA 32), middle frontal gyrus (BA 46/44), middle frontal gyrus (BA 46/9), superior frontal sulcus and lateral posterior nucleus of the thalamus. ROI coordinates are displayed in Table S1. The resulting variable served as the measure of WM network task activation.

### Statistical analysis

All other outcome variables used in analyses were checked for normality, transformations applied as necessary, and the models were refitted if there were deviations from the assumptions. All tests and confidence intervals for the primary hypotheses were two-sided and performed at the 0.05 level of significance. Our first hypothesis was that ketamine CMRO_2_ would be greater than saline CMRO_2_. To evaluate the difference between ketamine and saline, we performed a one-sample t-test on the relative percent signal change [(ketamine-saline)/saline], i.e., delta. If delta is significantly greater or less than zero, then the null hypothesis would be rejected. To amplify and complement these results, we also evaluated R_2_’, CBF, and CMRO_2_ in PFC and in SSM, VC, and GM.

Furthermore, we evaluated the association between task-related activation in PFC_NET (see above) and resting DLPFC CMRO_2_ under ketamine and under saline using mixed models. In these models, task-related activation in the WM network under saline or ketamine served as the endpoint. Load (2-target vs. 4-target) and phase (early retention vs. late retention) constituted the within-subject factors. Resting DLPFC CMRO_2_, evaluated in separate models, served as the continuous independent variable, and was centered for interpretability of all model effects. In all analyses, we fitted full models with all possible interactions, including those between the continuous independent variable and the factors. Unstructured variance-covariance matrix of the repeated measures was employed. We evaluated the association between ketamine-associated impairment in WM accuracy and resting CMRO_2_ under ketamine and saline in a similar manner. Ketamine-associated impairment in WM accuracy was quantified as ketamine percent correct minus saline percent correct. The models were considered exploratory and so were not corrected for multiple comparisons.

## Results

The median time between the first fMRI session and the second was 19 days. Ketamine plasma levels were 146 ng/dl ± 38.3 (range: 116-161 ng/dl). Ketamine reduced task-related activation and WM accuracy but not significantly. There was no significant relationship between plasma ketamine level and overall WM accuracy. Robust, statistically significant ketamine-associated increases in positive, negative, and dissociative symptoms were observed (Figure 1).

**Figure 1.**
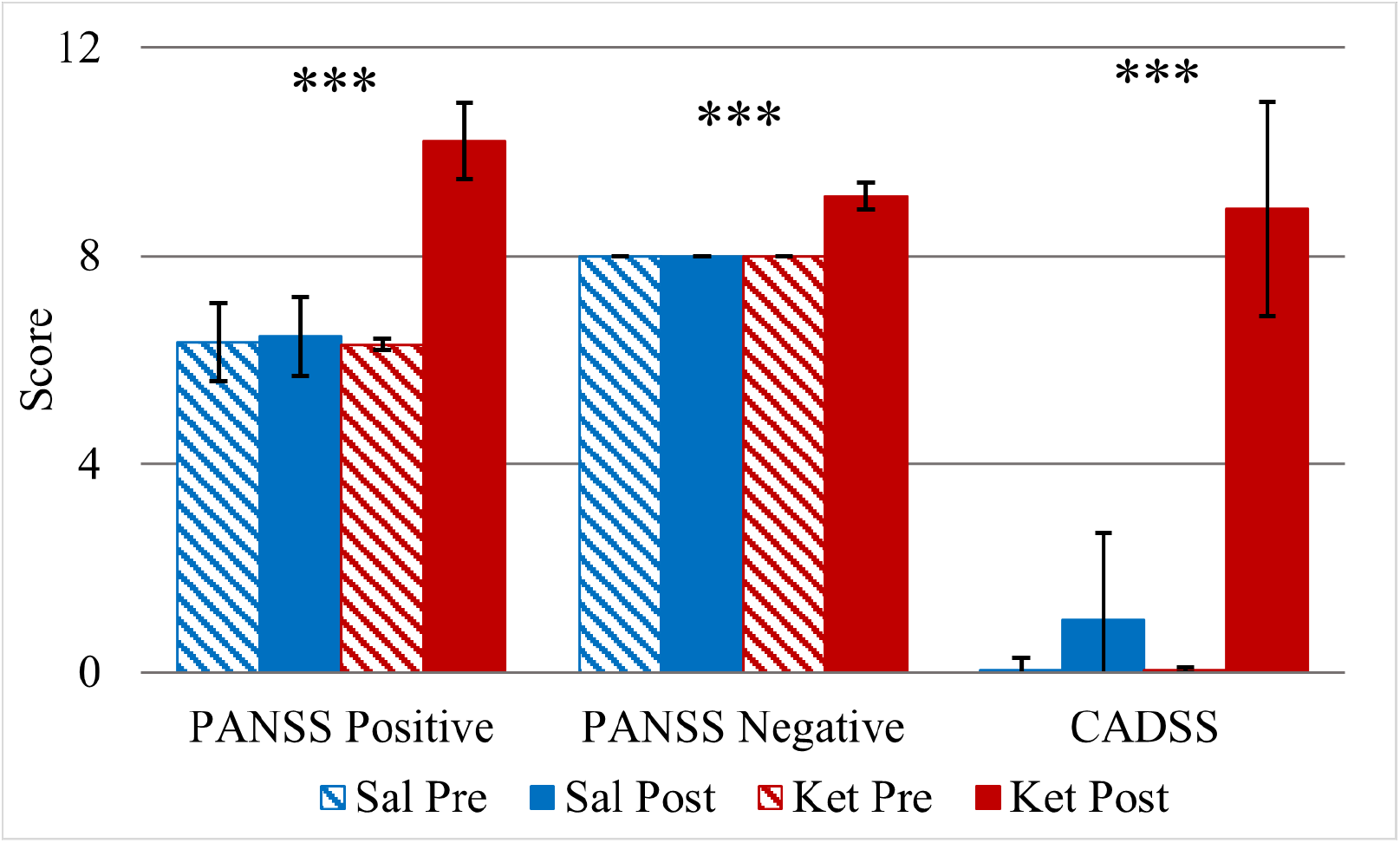
Factor scores for positive and negative symptoms on the Positive and Negative Syndrome Scale (PANSS) and total score on the Clinician-Administered Dissociative States Scale (CADSS) before scan (Pre) and after scan (Post) under saline (blue) and ketamine (red), Means ± SEM. *** = p < 0.001

### Ketamine increases CMRO_2_ in the prefrontal cortex

(Figures 2–5). There was no significant R_2_’ difference between the ketamine and saline conditions. Ketamine increased PFC CMRO_2_ by 15.6 ml/100g/min ± 4.86 SEM, t(22) = −3.21, p = 0.004. It produced a mean 13% increase ± 4.17 SEM in CMRO_2_, t(22) = 3.1 < 0.003. There was no statistically significant relationship between average plasma ketamine and the magnitude of the increase in CMRO_2_.

**Figure 2.**
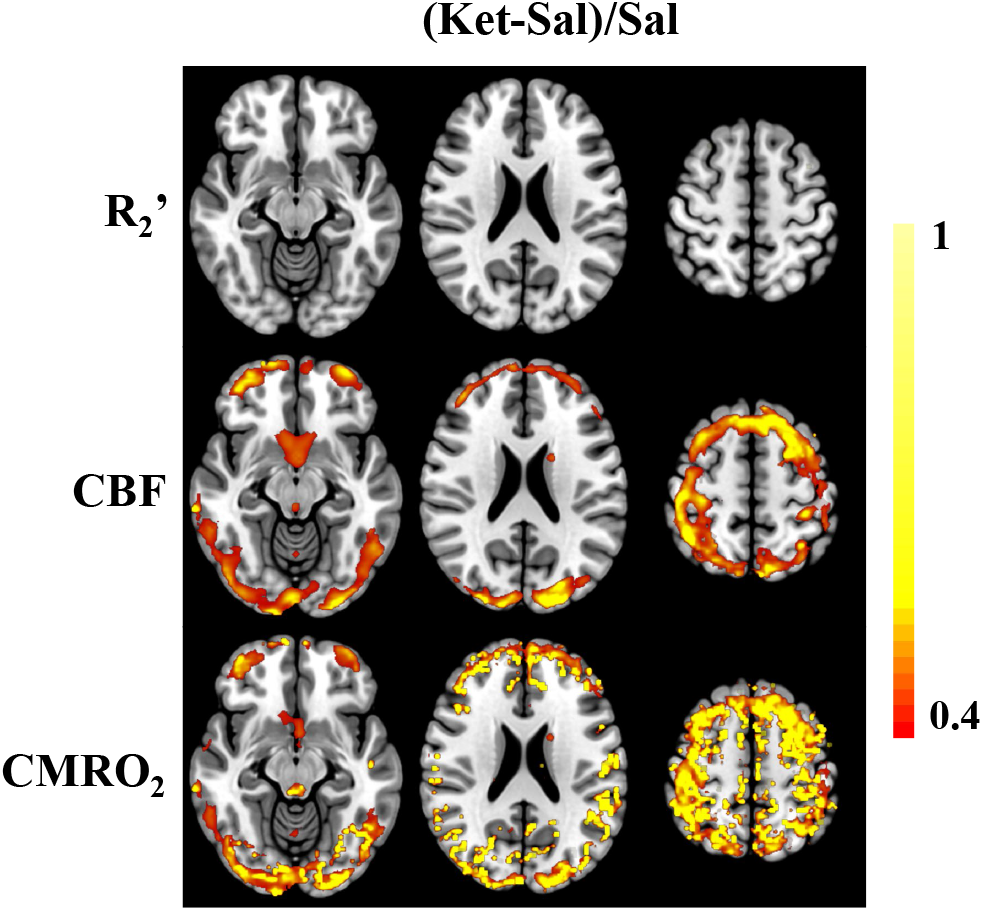
Relative percent signal change [(ketamine- saline)/saline] by modality. Cluster correction = 80.

PFC CBF correlated with PFC CMRO_2_ under saline, r(23) = 0.84, p < 0.001, and under ketamine, r(23) = 0.88, p < 0.001 (Tables 2 and 3). In addition, saline and ketamine PFC CBF were highly correlated, r(23) = 0.67, p < 0.001 (Table 4). Saline and ketamine PFC CMRO_2_ also correlated highly (r(23) = 0.73, p < 0.001 (Table 5). The findings above were paralleled in ROIs that were prespecified in our data analysis plan, i.e. SSM, VC, and GM (Figures 1–4, Tables 2–5). CBF and CMRO_2_ correlations during saline and during ketamine between ROIs were statistically significant and large (Table S2-S3). For example, CMRO_2_ correlations under saline between ROIs ranged from 0.67 to 0.92. All the inter-regional correlations were statistically significant. There was no statistically significant effect of ketamine on GBC or on GBC with signal removed (Figure 6).

**Figure 3.**
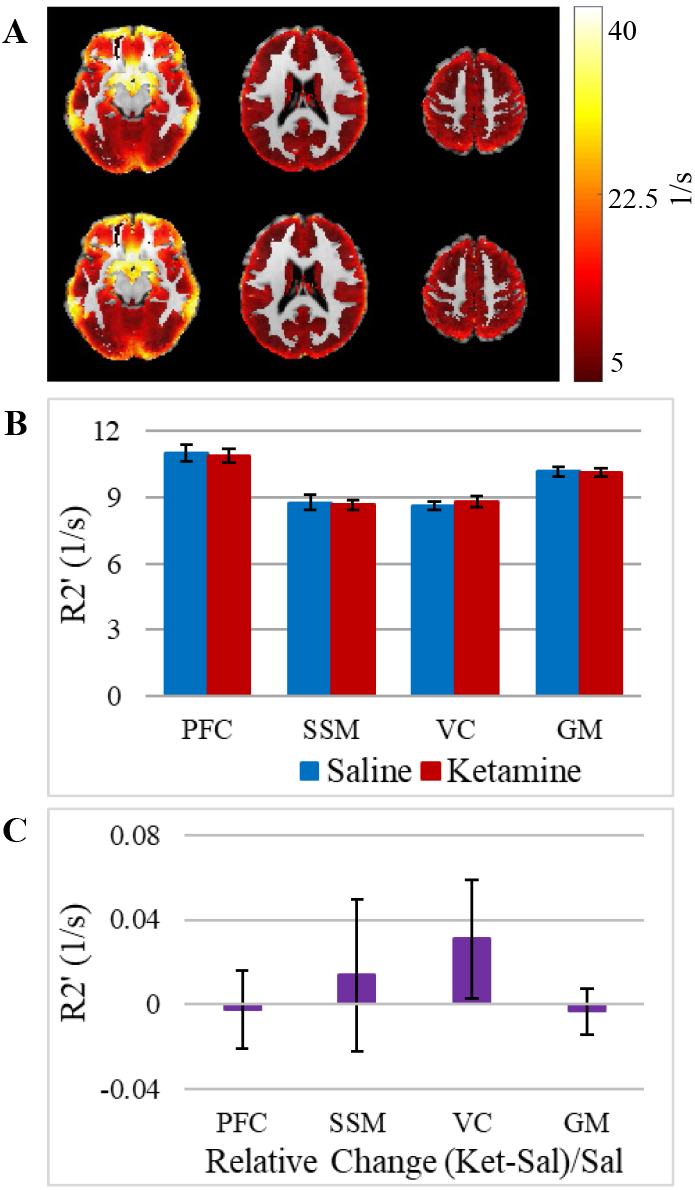
Ketamine does not significantly impact R_2_’. **A)** Average group R2’ maps, saline (top) and ketamine (bottom). **B)** R2’ under ketamine and saline in ROIs. **C)** Relative change in R2’ in ROIs.

**Figure 4.**
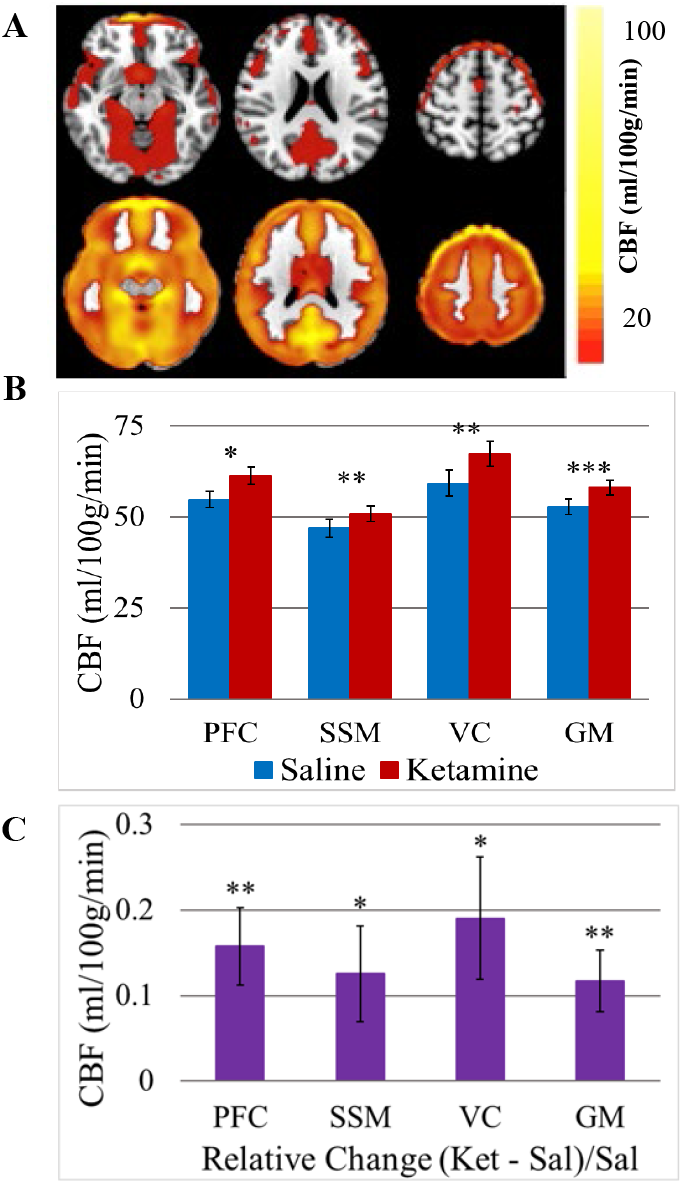
Ketamine increases cerebral blood flow (CBF). **A)** Average group CBF maps comparing saline (top) with ketamine (bottom). **B)** CBF under ketamine and saline in ROIs, Mean ± SEM. **C)** Relative change in CBF in ROIs, Mean ± SEM.* = p < 0.05, ** = p < 0.01, *** = p < 0.001

**Figure 5.**
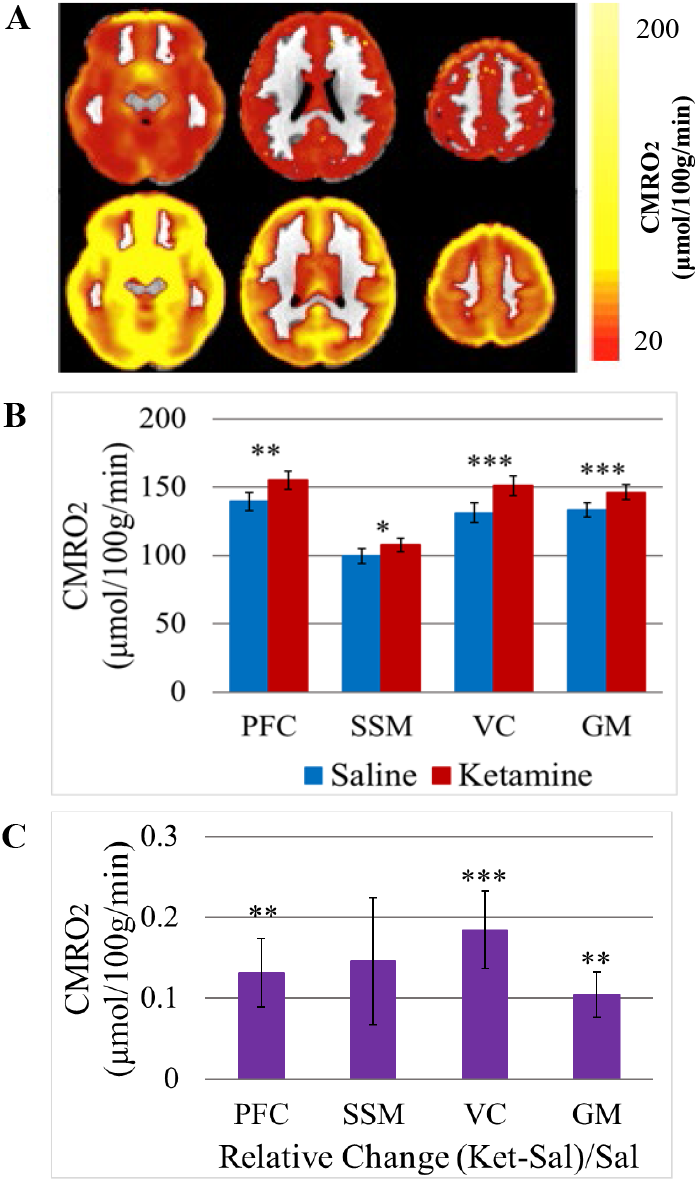
Ketamine increases cerebral metabolic rate of oxygen (CMRO_2_). **A)** Average CMRO_2_ maps under saline (top) and under ketamine (bottom). **B)** CMRO_2_ under ketamine and saline in ROIs, Means ± SEM. **C)** Relative change in CMRO_2_ in ROIs. * = p < 0.05, ** = p < 0.01, *** = p < 0.001

**Figure 6.**
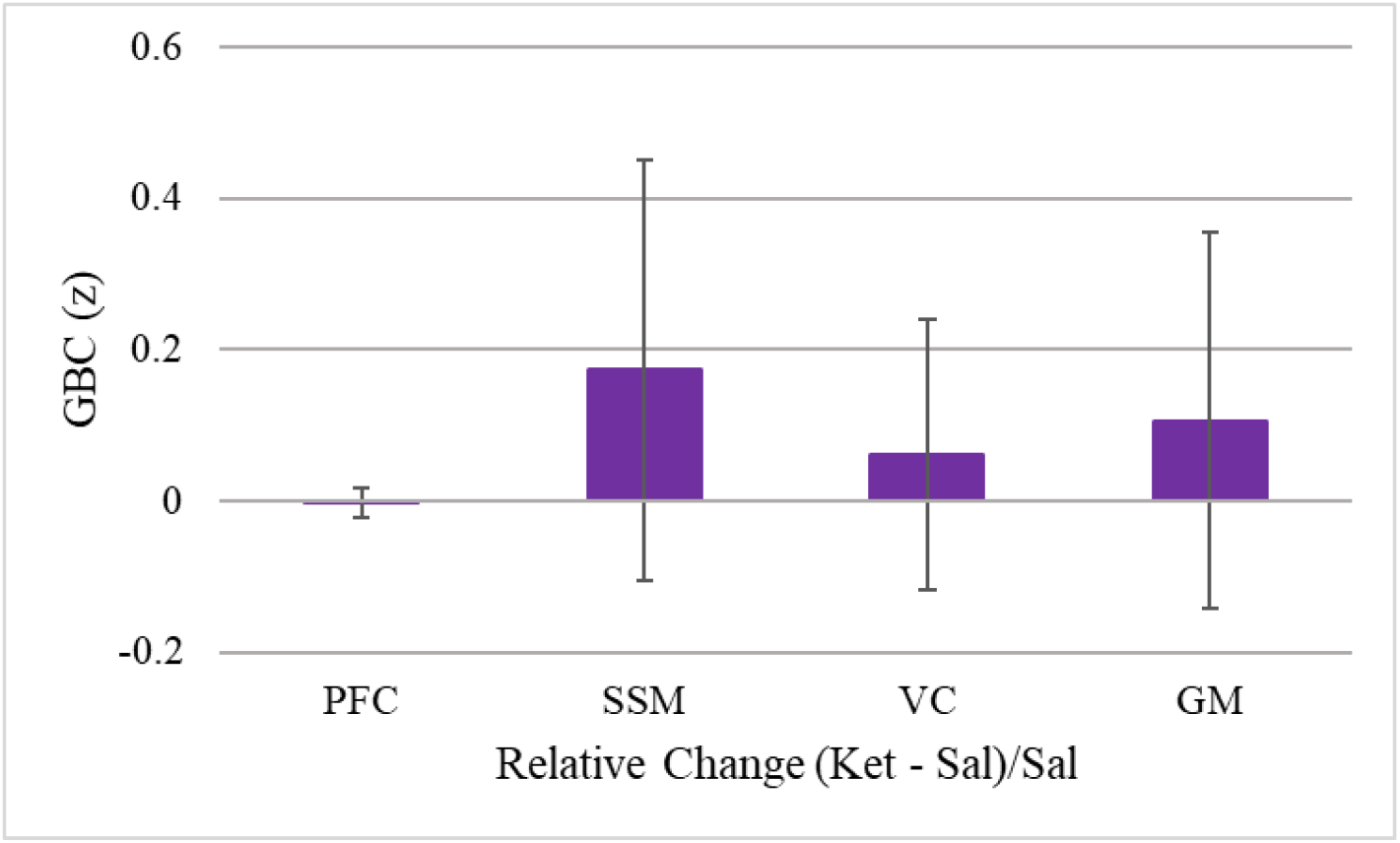
No ketamine-associated difference in GBC (global signal included).

**Table 2.** During Saline: High CBF-CMRO_2_ Correlations. * = Correlation is significant at the 0.05 level (2-tailed). ** = Correlation is significant at the 0.01 level (2-tailed).

|  |  | CMRO<sub>2</sub> PFC (Saline) | CMRO<sub>2</sub><br>Somatosensory and<br>Motor Cortex (Saline) | CMRO<sub>2</sub> Visual Cortex<br>(Saline) | CMRO<sub>2</sub> Grey Matter<br>(Saline) |
| --- | --- | --- | --- | --- | --- |
| CBF PFC (Saline) | Pearson Correlation | .835** | .684** | .533** | .722** |
|  | Sig. (2-tailed) | <.001 | <.001 | 0.009 | <.001 |
|  | N | 23 | 23 | 23 | 23 |
| CBF Somatosensory<br>and Motor Cortex<br>(Saline) | Pearson Correlation | .813** | .824** | .652** | .753** |
|  | Sig. (2-tailed) | <.001 | <.001 | <.001 | <.001 |
|  | N | 23 | 23 | 23 | 23 |
| CBF Visual Cortex<br>(Saline) | Pearson Correlation | .596** | .495* | .898** | .804** |
|  | Sig. (2-tailed) | 0.003 | 0.016 | <.001 | <.001 |
|  | N | 23 | 23 | 23 | 23 |
| CBF Grey Matter<br>(Saline) | Pearson Correlation | .827** | .684** | .781** | .887** |
|  | Sig. (2-tailed) | <.001 | <.001 | <.001 | <.001 |
|  | N | 23 | 23 | 23 | 23 |

**Table 3.** During Ketamine: High CBF-CMRO_2_ correlations. ** = Correlation is significant at the 0.01 level (2-tailed).

|  |  | CMRO<sub>2</sub> PFC (Ketamine) | CMRO<sub>2</sub> Somatosensory and Motor Cortex (Ketamine) | CMRO<sub>2</sub> Visual Cortex (Ketamine) | CMRO<sub>2</sub> Grey Matter (Ketamine) |
| --- | --- | --- | --- | --- | --- |
| CBF PFC (Ketamine) | Pearson Correlation | .877** | .855** | .567** | .821** |
|  | Sig. (2-tailed) | <.001 | <.001 | 0.005 | <.001 |
|  | N | 23 | 23 | 23 | 23 |
| CBF Somatosensory and Motor Cortex (Ketamine) | Pearson Correlation | .789** | .907** | .657** | .814** |
|  | Sig. (2-tailed) | <.001 | <.001 | <.001 | <.001 |
|  | N | 23 | 23 | 23 | 23 |
| CBF Visual Cortex (Ketamine) | Pearson Correlation | .698** | .659** | .905** | .860** |
|  | Sig. (2-tailed) | <.001 | <.001 | <.001 | <.001 |
|  | N | 23 | 23 | 23 | 23 |
| CBF Grey Matter (Ketamine) | Pearson Correlation | .858** | .845** | .770** | .918** |
|  | Sig. (2-tailed) | <.001 | <.001 | <.001 | <.001 |
|  | N | 23 | 23 | 23 | 23 |

**Table 4.** CBF correlations between saline and ketamine. * = Correlation is significant at the 0.05 level (2-tailed). ** = Correlation is significant at the 0.01 level (2-tailed).

|  |  | CBF PFC (Ketamine) | CBF Somatosensory and Motor Cortex (Ketamine) | CBF Visual Cortex (Ketamine) | CBF Grey Matter (Ketamine) |
| --- | --- | --- | --- | --- | --- |
| CBF PFC (Saline) | Pearson Correlation | .665** | .756** | .540** | .670** |
|  | Sig. (2-tailed) | 0.001 | <.001 | 0.008 | <.001 |
|  | N | 23 | 23 | 23 | 23 |
| CBF Somatosensory and Motor Cortex (Saline) | Pearson Correlation | .555** | .720** | .595** | .644** |
|  | Sig. (2-tailed) | 0.006 | <.001 | 0.003 | <.001 |
|  | N | 23 | 23 | 23 | 23 |
| CBF Visual Cortex (Saline) | Pearson Correlation | 0.353 | .470* | .803** | .612** |
|  | Sig. (2-tailed) | 0.098 | 0.023 | <.001 | 0.002 |
|  | N | 23 | 23 | 23 | 23 |
| CBF Grey Matter (Saline) | Pearson Correlation | .614** | .735** | .760** | .763** |
|  | Sig. (2-tailed) | 0.002 | <.001 | <.001 | <.001 |
|  | N | 23 | 23 | 23 | 23 |

**Table 5.** CMRO_2_ correlations between saline and ketamine. * = Correlation is significant at the 0.05 level (2-tailed). ** = Correlation is significant at the 0.01 level (2-tailed).

|  |  | CMRO<sub>2</sub> PFC (Ketamine) | CMRO<sub>2</sub> Somatosensory and Motor Cortex (Ketamine) | CMRO<sub>2</sub> Visual Cortex (Ketamine) | CMRO<sub>2</sub> Grey Matter (Ketamine) |
| --- | --- | --- | --- | --- | --- |
| CMRO<sub>2</sub> PFC (Saline) | Pearson Correlation | .732** | .806** | .782** | .789** |
|  | Sig. (2-tailed) | <.001 | <.001 | <.001 | <.001 |
|  | N | 23 | 23 | 23 | 23 |
| CMRO<sub>2</sub> Somatosensory and Motor Cortex (Saline) | Pearson Correlation | .426* | .773** | .600** | .517* |
|  | Sig. (2-tailed) | 0.043 | <.001 | 0.002 | 0.012 |
|  | N | 23 | 23 | 23 | 23 |
| CMRO<sub>2</sub> Visual Cortex (Saline) | Pearson Correlation | 0.398 | .490* | .839** | .600** |
|  | Sig. (2-tailed) | 0.060 | 0.018 | <.001 | 0.002 |
|  | N | 23 | 23 | 23 | 23 |
| CMRO<sub>2</sub> Grey Matter (Saline) | Pearson Correlation | .676** | .749** | .870** | .830** |
|  | Sig. (2-tailed) | <.001 | <.001 | <.001 | <.001 |
|  | N | 23 | 23 | 23 | 23 |

### Higher resting DLPFC CMRO_2_ predicted reduced WM-related activation and reduced WM accuracy during ketamine

In these analyses, we concentrated on DLPFC, a region within the PFC which has been strongly identified with working memory (2, 11, 75). CMRO_2_ in DLPFC correlated highly with CMRO_2_ in PFC during saline, r(23) = 0.92, p < 0.001, and under ketamine, r(23) = 0.73, p < 0.001. Saline CMRO_2_ DLPFC correlated moderately with ketamine CMRO_2_ DLPFC, r(23) = 0.62, p = 0.002.

Controlling for load (2-target vs. 4 target) and phase (early vs. later retention), DLPFC CMRO_2_ during ketamine infusion was not associated with WM network-related activation assessed during ketamine infusion (Figure 7A). Despite this negative finding, saline CMRO_2_ DLPFC was a significant predictor of network task activation, controlling for load and phase, [F (1, 21) = 6.75, p = 0.02 (Figure 7B)]. Notably, saline CMRO_2_ DLPFC was not associated with WM network-related activation assessed under saline.

**Figure 7.**
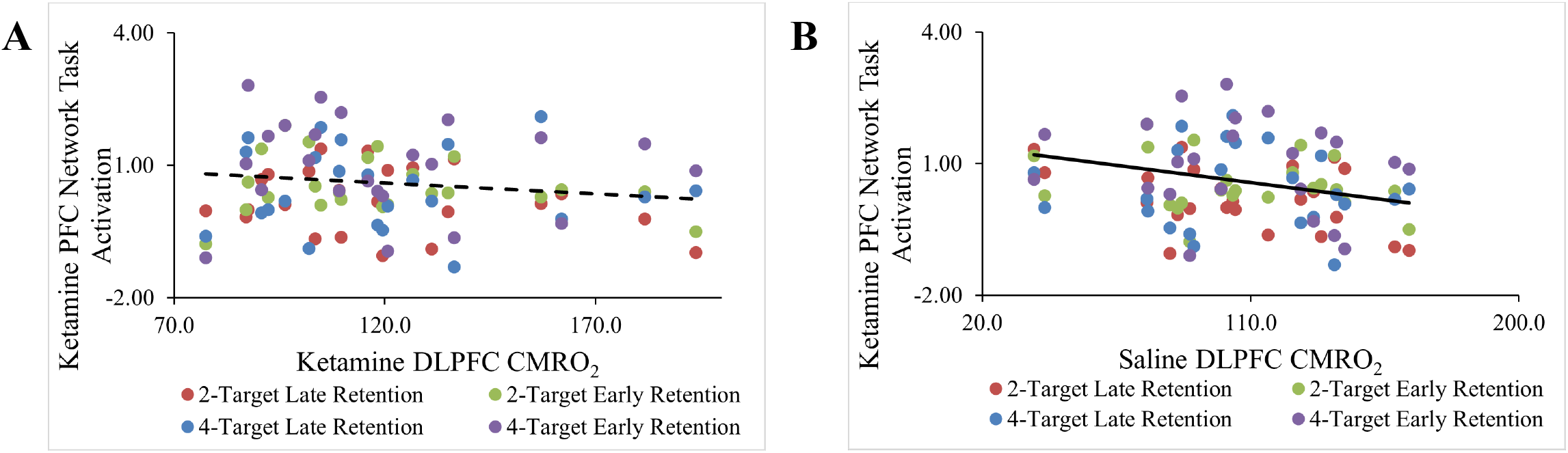
Participants with lower working memory task activation under ketamine had higher CMRO_2_ under saline and under ketamine. **A)** Non-significant relationship between Ketamine DLPFC CMRO_2_ and Ketamine Network Task Activation. **B)** Relationship between Saline DLPFC CMRO_2_ and Ketamine Network Task Activation, significance of the continuous independent variable (Saline DLPFC CMRO_2_), F (1,21) = 6.75, p = 0.02. One out-of-range value for a case during the 4-target retention interval not shown but included in the model.

Resting DLPFC CMRO_2_ during ketamine or saline predicted ketamine-associated impairments in WM accuracy. Controlling for load, higher ketamine CMRO_2_ was associated with greater ketamine-associated impairment in WM accuracy, F (1, 21) = 6.84, p = 0.02 (Figure 8A). Similarly, higher saline CMRO_2_ DLPFC was associated with greater ketamine-associated impairment, F (1, 21) = 5.37, p = 0.03 (Figure 8B). In all models relating CMRO_2_ to working memory activation or WM accuracy, there were no significant main effects of load or phase or interactions involving these factors on WM network related activation or WM accuracy during ketamine administration. Models using WM network task-related activation under saline as a dependent variable did not show relationships with CMRO_2_ assessed during ketamine or saline.

**Figure 8.**
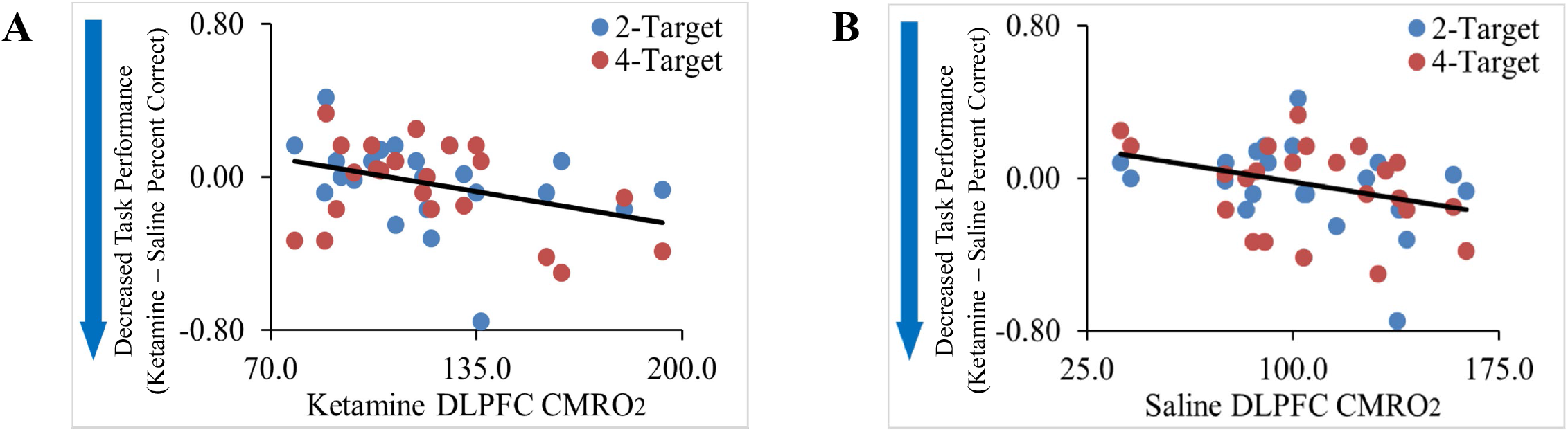
Participants with greater decreases in WM accuracy under ketamine had greater DLPFC CMRO_2_ under saline and under ketamine. **A)** Relationship between Ketamine DLPFC CMRO_2_ and ketamine-related impairments in working memory accuracy, significance of the continuous independent variable (Ketamine DLPFC CMRO_2_), F (1,21) = 5.37, p = 0.03. **B)** Relationship between Saline DLPFC CMRO_2_ and ketamine-related impairments in working memory accuracy, significance of the continuous independent variable (Saline DLPFC CMRO_2_), F (1,21) = 6.84, p = 0.02.

## Discussion

This study illustrated the use of calibrated fMRI to non-invasively measure drug effects on CMRO_2_. Using a gas-free calibrated fMRI technique, this study demonstrated that ketamine increased cortical CBF and CMRO_2_ and these two measures were highly correlated. R_2_’, a component of BOLD, was not sensitive to ketamine. Measurements of CBF and CMRO_2_ during saline were highly correlated with the respective measurements made during ketamine infusion. Reminiscent of the distribution of ketamine’s occupancy of NMDARs throughout the brain (76), the ketamine-related CBF and CMRO_2_ increases were widespread. CMRO_2_ under saline predicted WM network task activation during ketamine and ketamine-associated impairments in WM accuracy, even though the ketamine and saline infusions were performed a median of 19 days apart. Ketamine DLPFC CMRO_2_ did not predict ketamine WM network task-related activation, despite the fact that both measures were collected in the same scan day. However, it did predict the ketamine-associated WM accuracy deficit.

The study findings suggest that there may be an inverse relationship between resting cortical CMRO_2_ and WM-related PFC activation and WM performance. Higher CMRO_2_ during saline infusion predicted reduced WM-related PFC activation and reduced WM performance during ketamine. A link between these two conditions may be the functional status of various GABA interneuron populations. For example, in non-human primates, GABA_A_ receptor blockade was previously shown to both activate cortical neurons and to impair WM-related neural activity (29). Similarly, NMDAR antagonist administration was shown to reduce GABA neuronal activation, increase resting pyramidal neuronal activity in PFC and hippocampus, and impair WM accuracy (17). In animals, preventing the disinhibitory effects of ketamine via blockade of AMPA glutamate receptors or stimulation of metabotropic-2/3 receptors (mGluR2/3) reduces PFC hyperactivity and WM impairment (18).

In the current study, the magnitude of basal CMRO_2_ (saline CMRO_2_) was inversely related to WM-related cortical activation under ketamine. Effective WM task activation requires greater recurring excitation in response neurons in DLPFC Layer III corresponding to the stimulus relative to firing in other DLPFC neurons. Individuals who, at baseline, have higher basal metabolism may experience more homeostatic pressure under ketamine and have difficulty attaining task-related activation that is of the requisite magnitude for eidetic precision. Thus, the findings of this study may help to explain some of the inter-subject variability in response to ketamine. In addition, the inverse relationship between basal metabolism and WM task-related activation and WM accuracy, may help to explain why pretreatment with an mGluR2/3 agonist attenuates WM impairment in humans administered ketamine (77), which may have implications for treating symptoms (78) and cognitive impairments (30) associated with schizophrenia.

This study demonstrates tight CBF- CMRO_2_ coupling under ketamine, suggesting that neurovascular coupling remains intact under NMDAR blockade. Thus, this study strengthens confidence in the extant ketamine fMRI literature that employs traditional fMRI. Furthermore, it illustrates how calibrated fMRI can be used to assess neurovascular coupling in other drugs of interest such as anesthetics or psychedelics. Comparing baseline CBF and CMRO_2_ before and after drug may be a critical path to increasing confidence in neuropharmacological fMRI. Since inhalation is often anxiety-provoking, the non-gaseous technique described here has excellent potential for assessing resting baseline in a variety of drugs and in individuals with psychiatric disorders. Current theories discuss psychopathology as a disturbance in the balance of excitation and inhibition. Thus, a non-invasive way to characterize baseline differences would be extremely useful in testing and further articulating these theories.

The findings of the current study highlight that GBC and CMRO_2_ measure distinct features of cortical activation. The current study did not replicate our previous finding of overall functional connectivity increases under ketamine using the GBC technique, a data driven method used for studying functional connectivity (24). In this study, we used improved sequences and computed GBC with and without global signal removal. Another group has also reported negative overall findings (79), but has argued that components of the GBC signal derived using Principal Components Analysis reflect ketamine effects (80). Nonetheless, the lack of correlation between GBC and CMRO_2_ raises the possibility that these measures reflect distinct neuronal populations or distinct neuronal components, i.e., distal dendrites versus proximal elements (proximal dendrite, soma).

In this study, we were able to relate basal, resting CMRO_2_ to WM task activation and accuracy. These relationships were found despite a relatively small sample and the fact that the ketamine-associated reductions in WM task-related activation and in WM accuracy did not reach statistical significance. There is considerable inter-subject variability in ketamine-associated deficits in WM task-related activation and WM accuracy (81). Thus, non-replication could occur through chance in some samples. In addition, the other studies of WM and ketamine were single-blind and fixed order, whereas the current study was double-blind with order of saline and ketamine conditions randomized. This design may have eliminated some spurious sources of measured effects. A larger sample would allow us to test this more thoroughly and better sample individual responses to ketamine. Despite these issues, we were able to show, in an exploratory fashion, statistically significant relationships between resting saline or ketamine DLPFC CMRO_2_ and WM task activation and accuracy. The explanatory model discussed here would need to be prospectively tested in a larger sample.

In summary, this study combined saline and ketamine measurements of CBF and CMRO_2_ during rest with measurements of task-related activation and accuracy during a WM task. Potential functional connectivity changes under ketamine were assessed with a data-driven functional connectivity analysis technique, GBC. The analysis showed that ketamine increased CBF and CMRO_2_. Resting saline CMRO_2_ was significantly related to decreases in WM activation and WM accuracy, thus confirming hypotheses regarding NMDAR antagonism and WM. GBC with and without signal change was not sensitive to ketamine effects. This study highlights the importance of measuring baseline in using the tools of neuropharmacological fMRI and in constructing valid inferences regarding brain function in health and in pathology.

## Acknowledgements

This research was supported by the National Institute of Mental Health of the National Institutes of Health under Award Number 5R21MH110862-02 to NRD, JHK, FH; the U.S. Department of Veteran Affairs via its funding of the National Center for PTSD—Clinical Neurosciences Division (NRD, JHK); the Scottish Rite Schizophrenia Research Program to JHK; and the Yale Center for Clinical Investigation (CTSA Grant No. UL1 TR001863 from the National Center for Advancing Translational Science (NCATS), a component of the National Institutes of Health). The content is solely the responsibility of the authors and does not necessarily represent the official views of the National Institutes of Health or any other organization. The investigators acknowledge the critical role of the Yale Magnetic Resonance Research Center technicians and the Yale Center for Clinical Investigation nursing staff in completing this complex protocol.

