## Supplemental Information for "Ketamine Effects on Energy Metabolism, Functional Connectivity and Working Memory in Healthy Humans"

### List of Supplemental Data

**Supplemental Methods**

I. Calculation of $CMRO_2$.
II. Analysis of Resting-State Functional Connectivity (RSFC).

**Supplemental Figures**

- **Figure S1.** Regions-of-interest used in this study.

**Supplemental Tables**

- **Table S1.** Regions of interest used for assessing task-related activation during the spatial working memory task.
- **Table S2.** CBF inter-regional correlations under saline.
- **Table S3.** $CMRO_2$ correlations between saline and ketamine.

### Supplemental Methods

#### I. Calculation of $CMRO_2$

The basic equation for relating cerebral metabolism ($CMRO_2$) change between two conditions (C1 and C2) can be calculated from simultaneously measured changes in blood-oxygenation level dependent (BOLD) signal and cerebral blood flow (CBF) with the following equation:

$$\Delta BOLD/BOLD_{C1} = M\,[1 - (CMRO_{2,C2}/\,CMRO_{2,C1})^{\beta}\,(CBF_{C2}/CBF_{C1})^{\alpha-\beta}\,] \quad \text{[eq. 1]}$$

Parameters α and β are relatively stable and uniform through the brain and can be considered as region and condition independent (1-4). The constant β is an exponent linking the BOLD signal's dependencies on deoxyhemoglobin (deoxyHb) concentrations and relates $CMRO_2$ to CBF (2, 3). Parameter M is an echo-time (TE) dependent constant which incorporates the resting $CMRO_2$ and CBF conditions for a subject (2, 5). In eq. 1, M is the unknown parameter which must be independently measured in each subject so the $CMRO_2$ can be calculated.

Magnetic properties of blood change with blood oxygenation (or oxygen saturation),

$$Y = [oxyHb]/[deoxyHb+oxyHb], \quad \text{[eq. 2]}$$

and they affect the tissue water functional magnetic resonance imaging (fMRI) signal through intravoxel spin dephasing (6). This effect can be captured under well shimmed conditions by transverse relaxation rates measured by gradient-echo ($R_2$*) and spin-echo ($R_2$) (1-3), respectively:

$$R_2^* \cong R_2'(Y) + R_2(Y) + R_2(0) \quad \text{[eq. 2.1]}$$
$$R_2 \cong R_2(Y) + R_2(0) \quad \text{[eq. 2.2]}$$

where $R_2'(Y)$ and $R_2(Y)$ represent the BOLD relaxation components that are reversible (e.g., static magnetic fields inhomogeneity, slow diffusion regime) and irreversible (i.e., intermediate to fast diffusion regime), respectively (6), and $R_2(0)$ is the non-susceptibility-based effect (1, 2). Assuming that the intravascular weighting of these measured values is reduced by appropriate choice of TE and the non-susceptibility-based effect is negligible, the difference between the two rates gives:

$$R_2' = R_2^* - R_2 \quad \text{[eq. 2.3]}$$

where $R_2'$ represents the extravascular susceptibility-induced effects of tissue water. The parameter M obtained gas-free is simply given by the product of $R_2'$ and the TE (eq. 3). This M is the same that gas challenge experiments attempt to derive by measuring the maximum positive BOLD response (7), which occurs when all deoxyHb is assumed to be eliminated from vasculature with hypercapnia (8-11):

$$M = TE \cdot R_2' \quad \text{[eq. 3]}$$

Furthermore, based on the $R_2'$ equation we can simplify $CMRO_2$ calculation from eq. 1. The $R_2'$ value in a voxel is dependent on the CBV and the deoxyHb concentration ([deoxyHb]),

$$R_2' = A \cdot CBV \cdot [deoxyHb]^{\beta} \quad \text{[eq. 4.1]}$$

where A is a constant. Including the power law relationship between CBV and CBF (12) and between $CMRO_2$ and [deoxyHb] ($CMRO_2$ = [deoxyHb] · CBF),

$$R_2' = A \cdot c \cdot CBF^{\alpha} \cdot (CMRO_2 / CBF)^{\beta}, \quad \text{[eq. 4.2]}$$

which is

$$R_2' = A \cdot c \cdot CBF^{\alpha-\beta} \cdot CMRO_2^{\beta} \quad \text{[eq. 4.3]}$$

For changes of these parameters across the two conditions in the same session, constants A and k are eliminated, and it can be shown that:

$$(R_{2,C1}' / R_{2,C2}') = (CBF_{C1} / CBF_{C2})^{\alpha-\beta} \cdot (CMRO_{2,C1} / CMRO_{2,C2})^{\beta}, \quad \text{[eq. 5.1]}$$

and $CMRO_2$ changes (i.e., in each condition):

$$(CMRO_{2,C1} / CMRO_{2,C2}) = (R_{2,C1}' / R_{2,C2}')^{1/\beta} \cdot (CBF_{C1} / CBF_{C2})^{1-\alpha/\beta} \quad \text{[eq. 5.2]}$$

This equation shows that ratiometric measurements of $R_2'$ and CBF with values of $\alpha$ (0.1-0.3) and $\beta$ (1-1.5) detected within the physiological range can provide changes in $CMRO_2$.

### II. Analysis of Resting-State Functional Connectivity (RSFC)

The initial stages of the analysis were performed in the Center for Functional Magnetic Resonance Imaging of the Brain (FMRIB) Software Library (FSL) (13). High resolution multiband BOLD data was converted into Neuroimaging Informatics Technology Initiative (NIFTI) format and brain extraction performed to delete non-brain elements. Then images will be motion-corrected to the middle image of the run using MCFLIRT (13), corrected for slice-timing, smoothed with a Gaussian kernel of FWHM 5 mm and normalized with a single multiplicative factor. Voxel-wise temporal auto-correlation was estimated and corrected using the FMRIB's Improved Linear Model (FILM) (14). Images were low-pass filtered with a cut-off frequency of 0.08 Hz. Each individual's co-planar image was registered to their high-resolution structural scan and their high-resolution structural scan registered to the Montreal Neurological Institute (MNI) standard brain supplied with FSL.

To examine within-subject RSFC changes under ketamine, we produced whole-brain connectivity maps using the AFNI (http://afni.nimh.nih.gov/afni) 3dTcorrmap function, computing the correlation between each voxel and all the other voxels in the brain and returning an average correlation for each voxel. Global Brain Connectivity (GBC) results of subjects' resting runs were averaged. For comparison purposes, we ran the same analyses with global signal removed. To obtain the global signal, we computed the timecourse of all grey matter.

**Supplemental Figures**

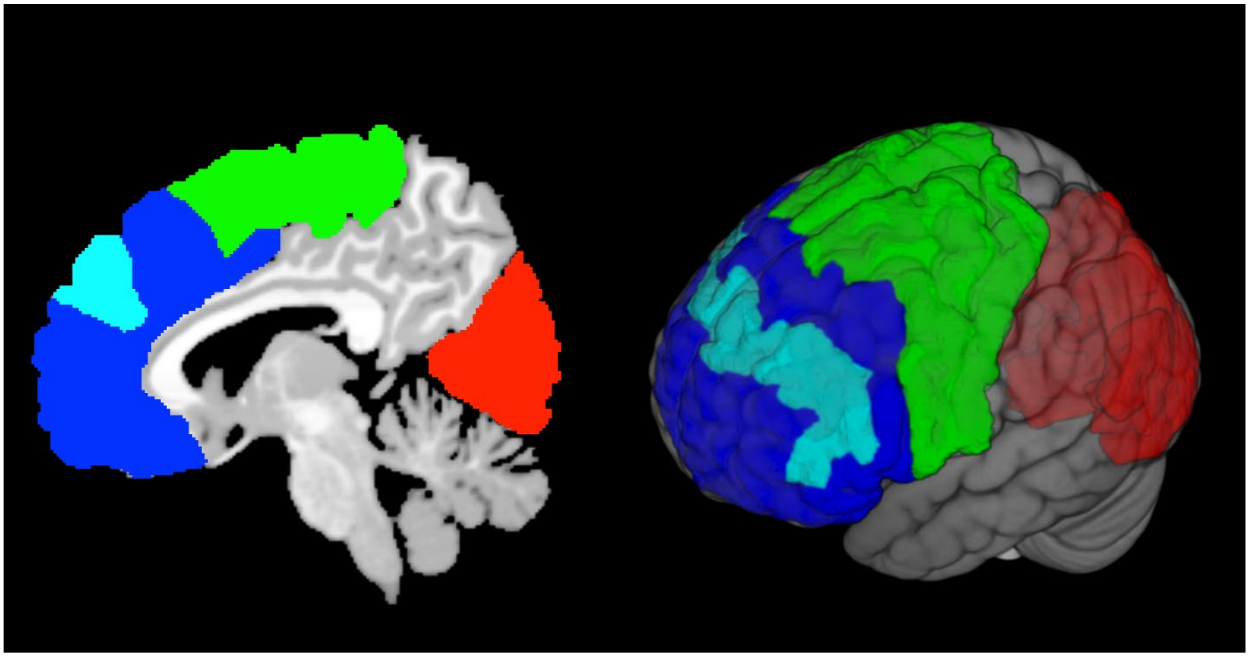

**Figure S1**. Regions-of-Interest used in this study. Red = Prefrontal Cortex, Green = Somatosensory and Motor Cortex, Dark Blue = Visual Cortex, Turquoise = DLPFC (within PFC).

**Supplemental Tables**

| Brodmann Area (BA) Name | BA Number | Right Hemisphere Coordinates | | |
|---|---|---|---|---|
| | | x | y | z |
| Inferior Frontal Gyrus | BA 45 | 37 | 19 | 9 |
| Anterior Corpus Callosum | BA 32 | 10 | 26 | 30 |
| Middle Frontal Gyrus | BA 46/44 | 48 | 30 | 20 |
| Middle Frontal Gyrus | BA 46/9 | 33 | 44 | 27 |
| Superior Frontal Sulcus | BA 8 | 24 | -2 | 50 |
| Lateral Posterior Nucleus of the Thalamus | n/a | 16 | -22 | 12 |

**Table S1.** Regions of interest used for assessing task-related activation during the spatial working memory task.

| | | CBF PFC (Saline) | CBF Somatosensory and Motor Cortex (Saline) | CBF Visual Cortex (Saline) | CBF Grey Matter (Saline) |
|---|---|---|---|---|---|
| CBF PFC (Saline) | Pearson Correlation | — | .876** | .627** | .899** |
| | Sig. (2-tailed) | | <.001 | 0.001 | <.001 |
| | N | 23 | 23 | 23 | 23 |
| CBF Somatosensory and Motor Cortex (Saline) | Pearson Correlation | | — | | |
| | Sig. (2-tailed) | | | | |
| | N | | 23 | 23 | |
| CBF Visual Cortex (Saline) | Pearson Correlation | | .639** | — | |
| | Sig. (2-tailed) | | 0.001 | | |
| | N | | 23 | 23 | |
| CBF Grey Matter (Saline) | Pearson Correlation | | .857** | .872** | — |
| | Sig. (2-tailed) | | <.001 | <.001 | |
| | N | | 23 | 23 | 23 |

**Table S2.** CBF inter-regional correlations under saline. ** = Correlation is significant at the 0.01 level (2-tailed).

| | | $CMRO_2$ PFC (Ketamine) | $CMRO_2$ Somatosensory and Motor Cortex (Ketamine) | $CMRO_2$ Visual Cortex (Ketamine) | $CMRO_2$ Grey Matter (Ketamine) |
|---|---|---|---|---|---|
| $CMRO_2$ PFC (Saline) | Pearson Correlation | — | .797** | .660** | .894** |
| | Sig. (2-tailed) | | <.001 | <.001 | <.001 |
| | N | 23 | 23 | 23 | 23 |
| $CMRO_2$ Somatosensory and Motor Cortex (Saline) | Pearson Correlation | | — | | |
| | Sig. (2-tailed) | | | | |
| | N | | | | |
| $CMRO_2$ Visual Cortex (Saline) | Pearson Correlation | | .667** | — | |
| | Sig. (2-tailed) | | <.001 | | |
| | N | | 23 | 23 | 23 |
| $CMRO_2$ Grey Matter (Saline) | Pearson Correlation | | .777** | .853** | — |
| | Sig. (2-tailed) | | <.001 | <.001 | |
| | N | | 23 | 23 | |

**Table S3.** CMRO2 correlations between saline and ketamine. ** = Correlation is significant at the 0.01 level (2-tailed).
